# Empirical validation of ephaptic coupling in printed human neural circuits

**DOI:** 10.1101/2025.05.21.655141

**Authors:** Johannes Striebel, Rouhollah Habibey, Daniel Wendland, Helge Gehring, Elizaveta Podoliak, Julia Sophie Pawlick, Kritika Sharma, Alex H. M. Ng, Wolfram Pernice, Volker Busskamp

**Affiliations:** University of Bonn, Faculty of Medicine, Department of Ophthalmology, Bonn, Germany; University Hospital Bonn; Institute of Physics and Center for Nanotechnology, University of Münster, Münster, Germany; Kirchhoff-Institute for Physics, University of Heidelberg, Heidelberg, Germany; Department of Genetics, Blavatnik Institute, Harvard Medical School, Boston, MA, USA and Wyss Institute for Biologically Inspired Engineering at Harvard University, Boston, MA, USA

## Abstract

Ephaptic coupling is a phenomenon describing the influence of endogenous electric fields on neuronal activity^1,2^. Although ephaptic coupling is deemed to contribute to computations in the brain^1,3,4^, the olfactory system^5^, the retina^6,7^, and to cardiac conduction^8,9^, and being associated with diseases like epilepsy^10,11^ and arrhythmia^12,13^, it is still poorly understood since it is notoriously difficult to investigate *in vivo* and *in vitro*. *In vitro* electrophysiology allows accessible and flexible experimentation, but circuits form randomly, leading to a lack of precision and reproducibility. Here, we present a method for reproducibly constructing human neuronal networks *in vitro* with single-cell precision. We constructed neuronal circuits from the bottom up by bringing their axons into close contact. This enabled us to measure the effects of ephaptic coupling and validate theoretical predictions, such as reduced action potential velocity, increased activity synchronization^14–16^ and reduced stimulation threshold^17^. Our precise measurements of electrophysiological activity support the importance of ephaptic coupling in neuronal circuit function. Printed neuronal circuits allow detailed *in vitro* studies of neuronal interactions and may serve as a platform for disease modeling related to synaptic, ephaptic, or myelination processes.

## Introduction

Ephaptic coupling is a mechanism of neuronal interaction in addition to chemical and electrical synapses. Electric fields present in neuronal tissue due to the flow of ions and charged neuronal compartments have been shown to affect functional properties such as signal propagation or synchronization in surrounding tissue. For example, propagation speed and synchronization were reported in crab nerves^16^. Because ephaptic coupling is mediated by electric fields, it is not easily manipulated experimentally and is therefore difficult to study. While computational studies have extensively explored the potential consequences of ephaptic coupling^3,15,17–23^, there were some experimental studies that have had to use external fields or advanced methods to study ephaptic effects ^4,14,24,25^. However, its role in neural computation and its functional significance in the brain are still open questions in need of further experimental validation. There is evidence that ephaptic coupling affects olfaction^5^, the retina^6,7^, and possibly higher-level computations of the brain^1,3,4^. Cardiac conduction may also be supported by ephaptic mechanisms in addition to gap junctions^26^. It is suggested that ephaptic coupling may also contribute to pathological conditions such as epileptic seizures^10,11,20,21,27^ and arrhythmias^12,13,18,19^.

We reasoned that to better understand this phenomenon, we would need to create an experimental platform that allows precise control over individual human induced pluripotent stem cell (hiPSC)-derived neurons *in vitro*. Until now, *in vitro* networks have suffered from several drawbacks. First, most experiments have a large inter-batch variability in network architecture. This can be partially addressed by using mechanical confinements with predefined structures such as microfluidics or surface patterning with chemical cues ^28–32^, but reproducible connectivity down to the single cell level has not yet been achieved. Second, uncontrolled growth and network structure contribute to the complexity of data acquisition and analysis. Third, there is variation in network morphology over time, making it difficult to study them longitudinally^33^. To overcome these problems, we have developed a method to build neuronal circuits from the bottom up with single-cell resolution. We call this Single-neuron Network Assembly Platform (SNAP). To the best of our knowledge, it has not previously been possible to reproduce defined neural circuits with single-cell precision.

## Results

### SNAPs react to synaptic intervention

To build SNAPs, we chose to use microfluidics as a scaffold in which human neurons can grow and be guided to connect in a predefined manner. This is a well-established method for engineering neural circuits^34^ and has several advantages over other methods. First, microfluidics interface well with microelectrode arrays, which facilitate electrophysiological readout of neuronal activity over the lifetime of the circuit. Second, developing neural networks *in vitro* show substantial changes in morphology over time^33^. Microfluidics keeps the circuit structure constant over time and the neurites close to the readout electrodes. Third, the stable microenvironment within microfluidic devices can shield cells from mechanical perturbations, increasing the survival of neuronal circuits during media changes or recording sessions. Alternative approaches like microcontact printing of adhesive molecules showed unspecific growth and unwanted connections as well as morphological instability over time^35^. We used the transgenic human inducible Neurogenin (iNGN) cell line in which the overexpression of the transcription factors Neurogenin-1 and -2 drives hiPSCs into postmitotic neurons in just four days^36^.

Simply pipetting neurons onto the microfluidic scaffolds did not ensure that each microwell contained exactly one cell. To build the circuits in a tightly controlled, bottom-up fashion, and to incorporate multiple cell types in a reproducible manner, we needed a method that would allow precise placement of individual cells in each microwell (Fig. 2a). We were inspired by blastocyst injection, a technique used to generate transgenic mouse lines by transferring stem cells with a micropipette. To implement this, we developed a custom setup that included a microscope with a micromanipulator holding a glass pipette connected to a microinjector (Supplementary Fig. 1b, Supplementary Video 1). A critical aspect of the setup was to match the tip diameter of the micropipette to the cell dimensions to ensure efficient and reliable cell transfer.

**Fig. 1:**
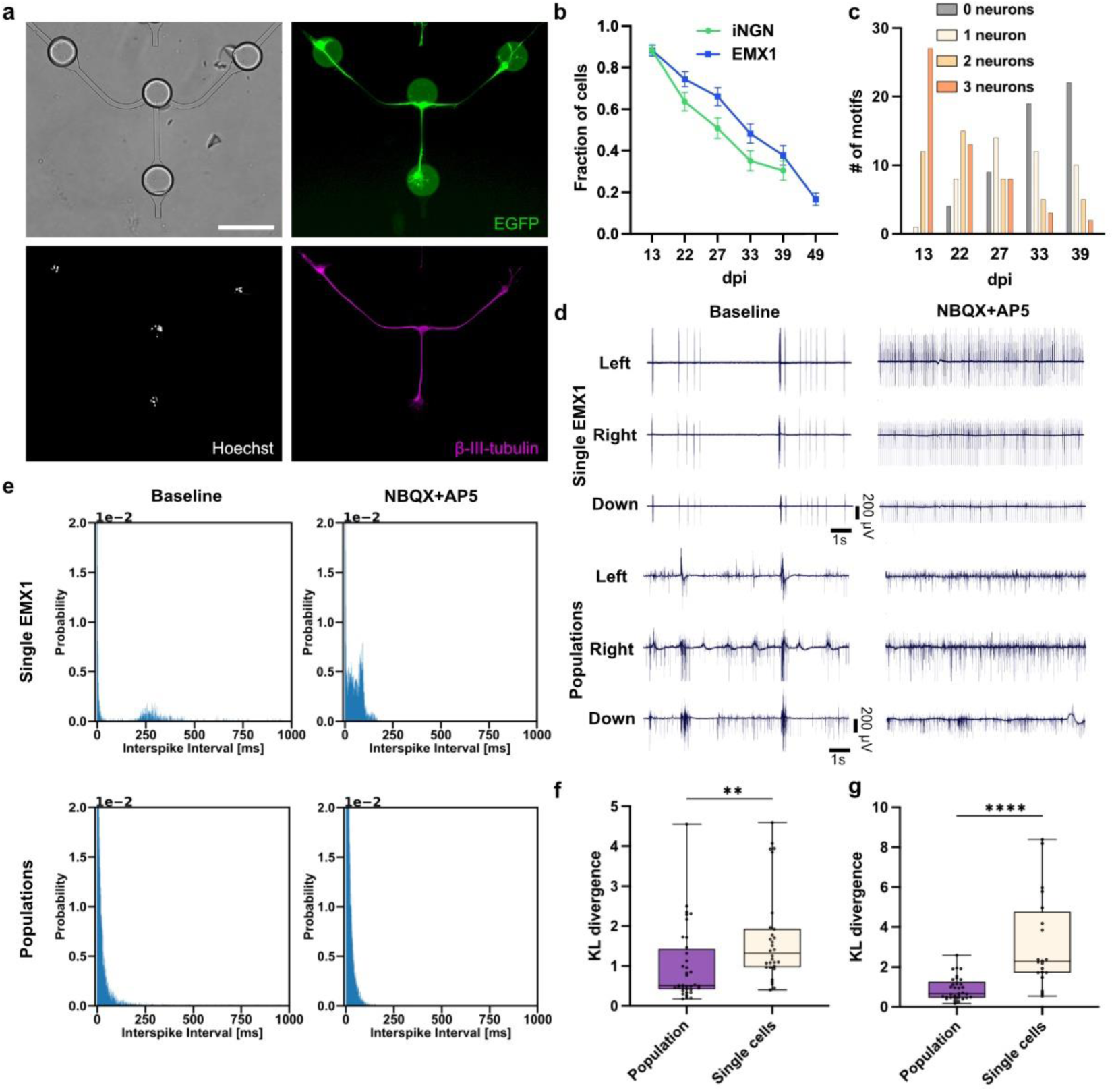
SNAPs are sensitive to synaptic antagonists. **a**, Representative neuronal circuit motif consisting of 3 iNGN neurons in microfluidics. Circular structures in the brightfield image are microwells with single neurons placed in outer wells. Immunofluorescence images of β-III-tubulin, Hoechst, and constitutively expressed EGFP. Scale bar, 200 µm. **b**, Fraction of surviving iNGN and EMX1 neurons in 3-neuron circuit motifs over time at different days post induction (dpi) (mean±s.e.m.; iNGN: n = 64 circuits; EMX1: n = 60 circuits). **c**, Number of circuit motifs with respective number of iNGN neurons per dpi. 10 samples with 4 motifs each were monitored. **d**, Representative raw traces of three electrodes (left, right, down channels) in one circuit motif of a SNAP (3x EMX1 neurons) and populations of EMX1 neurons seeded in the same structure. Left, baseline recordings. Right, with excitatory blockers NBQX+AP5. **e**, Interspike interval (ISI) histograms of the activity measured in representative electrodes of a circuit motifs with single and population EMX1 neurons. Left, baseline recordings. Right, after adding excitatory blockers NBQX+AP5. For the single cell case a change in histogram shape is visible. **f**,**g**, Kullback-Leibler (KL) divergence of baseline vs. NBQX+AP5 ISI histograms for single neuron and population circuits. Both EMX1 (**f**) and iNGN (**g**) neurons show a statistically significant increase in KL divergence for the single neuron circuits compared to populations, indicating a higher sensitivity to interventions via synaptic antagonists. In both plots, Box: 25^th^ to 75^th^ percentile with median. Whiskers min to max. Two-tailed Mann-Whitney test (EMX1: n = 36 and 30 electrodes for population and single cell respectively; iNGN: n = 33 and 20 electrodes for population and single cell respectively), ** *P≤0.01*, **** *P≤0.0001*.

**Fig. 2:**
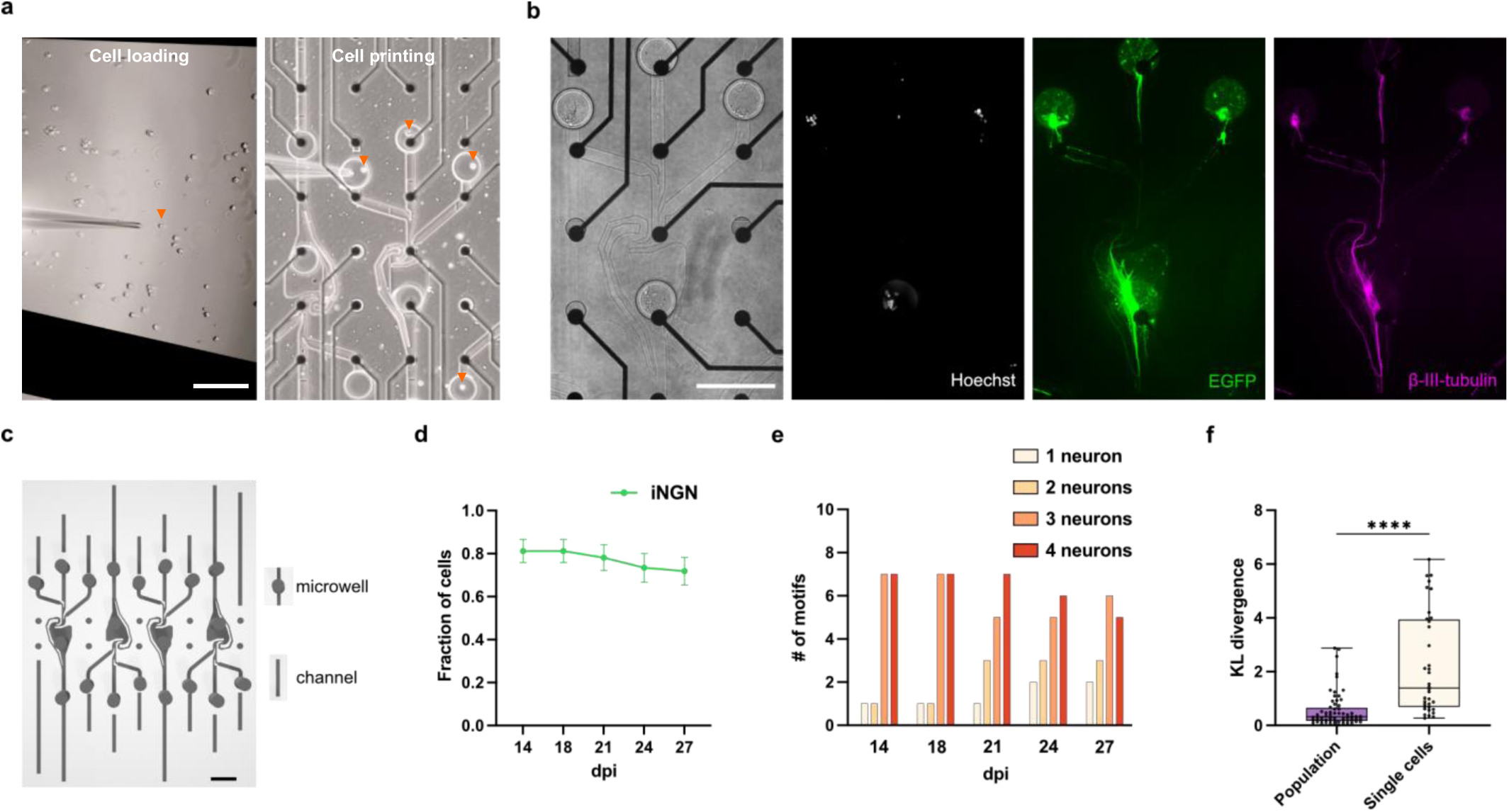
Directional neuronal circuits with single neuron resolution. **a**, Left, single cell (orange arrow) being taken up by micropipette. Right, cell release from micropipette into microwell. All single cells placed in microwells are marked with an orange arrow. Microfluidic structure is aligned to recording electrodes (black dots) of a multielectrode array (MEA). **b**, Representative neuronal circuit motif consisting of 4 iNGN neurons in microfluidics. Circular structures in the brightfield image are microwells with single cells placed in all wells. The stomach-shaped structure in the middle guides neurites downwards^41,42^. Immunofluorescence images of β-III-tubulin, Hoechst, and constitutively expressed EGFP. Scale bar, 200 µm. **c**, An array of four identical microstructures was designed to fit on one MEA. **d**, Fraction of viable cells at several days post induction (dpi) for directional 4-neuron circuit motifs as shown in (**b**) (mean±s.e.m., iNGN: n = 16 circuits). **e**, Number of circuit motifs containing respective number of iNGN neurons per dpi. Four samples with 4 motifs each were monitored. **f**, Kullback-Leibler (KL) divergence of baseline vs. NBQX+AP5 ISI histograms for directional SNAPs and population control. Single cell directional circuits show a statistically significant increase in KL divergence compared to populations of iNGNs in the same microfluidic structure, confirming functional excitatory synapses and indicating a higher sensitivity to interventions via synaptic antagonists. Box plots: 25^th^ to 75^th^ percentile with median. Whiskers: min to max. Two-tailed Mann-Whitney test (n = 64 and 36 electrodes for population and single cell respectively). **** *P≤0.0001*.

The next challenges were biocompatibility and neuronal survival as single cells per well. In stem cells, the ROCK pathway is commonly inhibited after passaging to avoid anoikis^37^. Recently, a more potent four-substance cocktail (CEPT) has been reported for this purpose, building on the idea of ROCK pathway inhibition^38^. We hypothesized that these supplements might help the initial survival of cells until a circuit is established. Indeed, the addition of a ROCK pathway inhibitor (ROCKi) improved survival up to 5 days after seeding, but showed no significant improvement thereafter (Supplementary Fig. 1c). In contrast, the addition of CEPT dramatically improved cell survival. Since iNGN neurons typically show robust electrophysiological activity by 14 days post induction (dpi), with functional synapses present by 21 dpi^39^, we had a method that allowed us to construct identical circuits and record functional activity from them in a manageable batch size.

We also created SNAPs composed of two different neuronal cell types, iNGN and iNGN cells in which the third transcription factor EMX1 was overexpressed (hereafter EMX1 cells, see Supplementary Fig. 1d).

First, we fabricated circuits consisting of 3 neurons in a Y-shaped microfluidic structure (Fig. 1a). Immunocytochemistry confirmed the structural integrity and neuronal identity of the cells. EMX1 neurons show a slightly better survival rate over time than iNGN neurons (Fig. 1b). Out of 40 circuits, more than 10 complete circuits were available for experiments after 22 dpi in the case of iNGN cells (Fig. 1c). To verify that the SNAPs were functionally connected via excitatory synapses, we added a mixture of the AMPA receptor antagonist (NBQX) and NMDA receptor antagonist (AP5) to the cultures. We observed a significant change in the electrophysiological activity when comparing the baseline activity with the antagonists added to the SNAPs for both iNGN and EMX1 neurons. As a control, we used cultures in which populations of neurons were added to the microwells instead of single neurons. There was also a change in the raw activity traces in the controls, although not as pronounced (Fig. 1d). Interspike interval (ISI) histograms are a fundamental measure of electrophysiological activity. For SNAPs, the ISI histograms were significantly changed after the addition of the antagonists, while this change was subtle for the population controls (Fig. 1e). Using the Kullback-Leibler (KL) divergence^40^, we were able to quantify how much the two histograms (baseline vs. NBQX+AP5) differ. Both iNGNs and EMX1 showed a significant increase in KL divergence for SNAPs compared to population controls (Fig. 1f,g), indicating that SNAPs provide a more sensitive readout to interventions than populations of cells. This could be due to the dense activity in neuronal populations masking such effects.

### Ephaptic coupling in SNAPs

For ephaptic coupling to be effective, it is important to bring axons into close contact. Therefore, we created specific microfluidic structures that unidirectionally guide axons of four neurons together^41^ (Fig. 2a-c). One multielectrode array (MEA) with an 8x8 array of electrodes accommodated a microfluidic device with four identical structures (Fig. 2c). Notably, the survival of neurons in these axon-guided structures was increased compared to the Y-shaped structure (Fig. 2d, e). We also tested that the circuits were responsive to synaptic antagonists to block transmission other than ephaptic coupling (Fig. 2f). Similar changes in electrophysiological activity and ISI histograms could be observed as quantified by the KL divergence. Cross-correlation analysis of the activity supported the directional signal propagation in the microfluidic structures (Supplementary Fig. 2c).

Next, we focused on ephaptic coupling characteristics such as changes in action potential (AP) velocity. Within SNAPs, we were able to precisely measure AP velocity changes in correlation to the number of different neuronal circuit members. We observed a significant decrease in AP velocity as more neurons interacted (Fig. 3a). To confirm that this effect was not mediated by synaptic delays, we measured AP velocity under excitatory synaptic antagonists NBQX+AP5 and electrical synaptic antagonist carbenoxolone (CBX), which did not change significantly (Fig. 3b). Our microfluidic structures have regions where axons are still separated and maintained individually and are later brought into contact to bundle. If ephaptic coupling is involved, we would have expected the AP velocity to be different in the areas of individual axons compared to the bundled areas. Indeed, we observed a significant decrease in AP velocity between these regions (Fig. 3c). In particular, our experimental setup facilitated a detailed study of individual circuits longitudinally over time. Figure 3d shows the evolution of AP velocity for different dpi in two circuits. Some neurons degenerate and are removed from the circuits over time, shifting the presence from bundled axons to individual axons. This resulted in a significant increase in AP velocity, further indicating that the change in cell number affected the propagation of APs. We observed a direct influence of axonal interaction length on AP velocity as predicted by theory^17^ (Fig. 3e). Immediately after seeding, we observed that axons were mostly separate, but over time they began to bundle together, resulting in a decrease in AP velocity. This suggests that the increased interaction length resulting from axon bundling influenced AP propagation. After degeneration of two of the four neurons in the circuit, AP velocity increased again, suggesting that the reduction in the number of neurons may have restored more efficient propagation. This change highlights the complex relationship between axonal organization and the dynamics of AP conduction in the circuit.

**Fig. 3:**
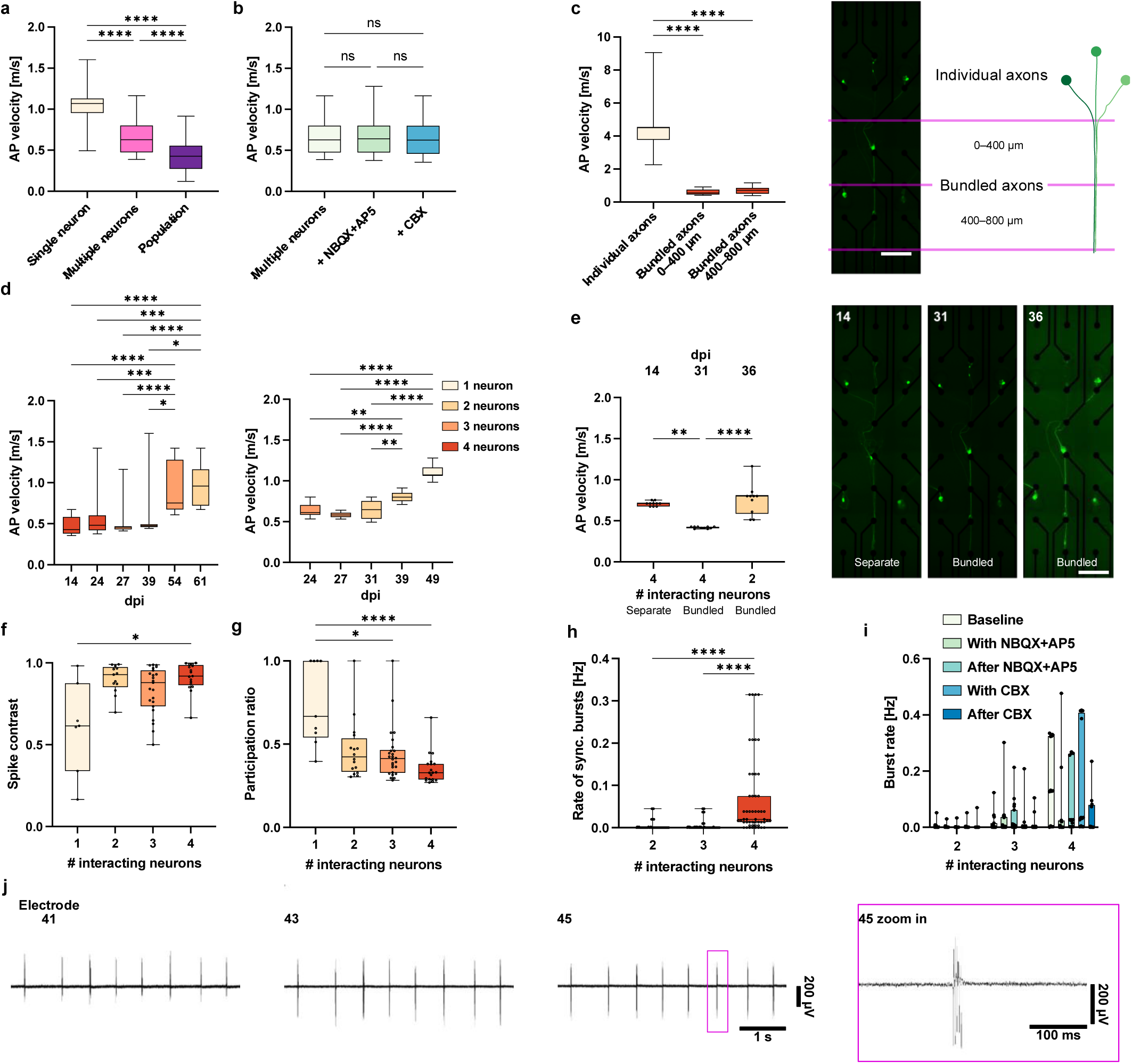
Ephaptic coupling in directional single neuron circuits. **a**, Velocity of action potential (AP) propagation for single iNGN neurons, multiple (2–4) neurons and neuronal populations in directional microfluidic structures. AP velocity decreases significantly with the number of neurons. Kruskal-Wallis test with Dunn’s multiple comparisons test (Single neurons: n = 60 APs; Multiple neurons: n = 100 APs; Population: n = 79 APs). **b**, AP velocity does not differ significantly for multiple neurons after adding excitatory blockers NBQX+AP5 or gap junction blocker CBX. Two-tailed Mann-Whitney test (Multiple neurons: n = 100 APs; Multiple neurons + NBQX+AP5: n = 100 APs; Multiple neurons + CBX: n = 100 APs). **c**, AP velocity differs significantly in areas of individual vs. bundled axons. Kruskal-Wallis test with Dunn’s multiple comparisons test (Individual axons: n = 29 APs; Bundled axons 0–400 µm: n = 50 APs; Bundled axons 400–800 µm: n = 50 APs). **d**, Development of AP velocities over time for 2 directional circuit motifs. Velocity increases significantly in the area of bundled axons with number of cells reducing in the circuit by natural cell death. Kruskal-Wallis test with Dunn’s multiple comparisons test (Left: n = 20, 20, 20, 19, 19, and 20 APs at dpi 14–61; Right: n = 20, 20, 22, 20, and 19 APs at dpi 24–49). **e**, Change of AP velocity in a circuit motif over time. At 14 dpi, axons are mostly separate leading to a short interaction length and higher velocity as compared to 31 dpi when axons have bundled together. After 2 cells in the circuit degenerated the velocity increased again. Kruskal-Wallis test with Dunn’s multiple comparisons test (n = 10 APs for each condition). **f**, Spike contrast as a measure of synchronized activity in the circuits is increasing significantly with the number of interacting neurons in directional circuits. Kruskal-Wallis test with Dunn’s multiple comparisons test (n = 7, 13, 23, and 17 circuits for 1–4 neurons respectively) after outlier identification with ROUT method (Q=1%). **g**, Participation ratio (PR) is decreasing with the number of neurons. A decrease in PR indicates a lower dimensionality and thus more coordinated neuronal network activity. Kruskal-Wallis test with Dunn’s multiple comparisons test (n = 9, 16, 26, and 19 circuits for 1–4 neurons respectively). **h**, Rate of synchronized burst activity in the circuit increases significantly with the number of neurons. Kruskal-Wallis test with Dunn’s multiple comparisons test (n = 48, 79, and 52 circuits for 2–4 neurons respectively). **i**, Burst rates in circuits increase with the number of neurons, irrespective of the presence of excitatory (NBQX+AP5) or gap junction (CBX) blockers (n = 11, 19, and 12 circuits for each condition with 2–4 neurons respectively for each condition). J, Exemplary raw traces measured in the area of bundled axons of one directional circuit. A strong synchronization of activity is visible. In all plots, ns *P>0.05*, * *P≤0.05*, ** *P≤0.01*, *** *P≤0.001*, **** *P≤0.0001.* Box plots: 25^th^ to 75^th^ percentile with median. Whiskers: min to max. Scale bars, 200 µm.

Another hallmark of ephaptic coupling is synchronization^14–16^. The raw signals of directional SNAPs already showed synchronized activity (Fig. 3j). Spikes occurred mostly in dense bursts of APs preceded and followed by longer periods of no spikes. This notion was supported by the fact that spike-contrast, a measure of synchrony, increased with the number of interacting neurons^43^ (Fig. 3f). The participation ratio^44^, a measure of the dimensionality of network activity, also decreased with the number of interacting neurons, further indicating increased synchronicity (Fig. 3g). Other classical measures of synchronicity such as burst rate and rate of synchronous network bursts also increased with number of interacting neurons (Fig. 3h,i).

### Stimulation of SNAPs

Ephaptic coupling is also predicted to lower the stimulation threshold of neurons within a circuit^23^, meaning that electrical stimuli applied to networks with more cells should elicit stronger responses. Indeed, we found that firing rates under stimulation increased superlinearly with the number of interacting neurons (Fig. 4a). This was also observed when excitatory synaptic antagonists were applied. In addition to electrical stimulation, we also used optogenetic stimulation to target single iNGN neurons expressing fChRimson-EYFP^45^, and measured their activity using non-invasive multielectrode array technology. Light stimulation elicited spikes from single neurons. As a further control for our ephaptic coupling experiments, we measured the AP velocity in the optogenetically activated neurons and found them to be comparable to spontaneously active neurons, with no decline in AP propagation velocity (Fig. 4b).

**Fig. 4:**
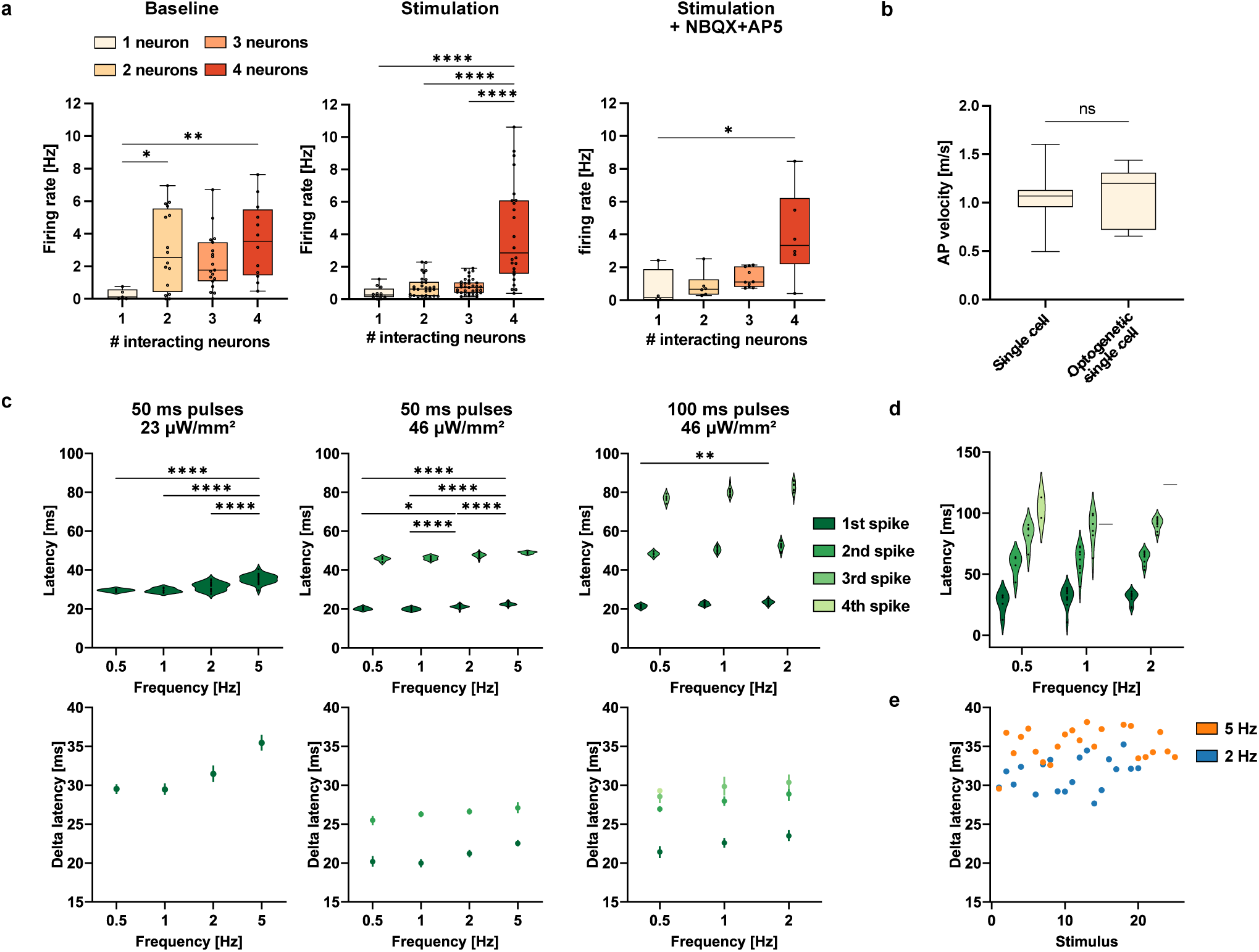
SNAPs can be stimulated electrically and optogenetically. a, Firing rates in directional SNAP circuits without stimulation (baseline) and while electrically stimulating one electrode in the circuit with and without excitatory blockers NBQX+AP5. Rates increase exponentially with the number of neurons under stimulation irrespective of blockers being present. Kruskal-Wallis test with Dunn’s multiple comparisons test (Firing rates measured in individual electrodes of 16 circuits included in Baseline: n = 5, 16, 19, and 12; Stimulation: n = 12, 30, 37, and 24; Stimulation + NBQX+AP5: n = 4, 6, 10, and 6 for 1–4 neurons respectively) after outlier identification with ROUT method (Q = 1%). **b**, Action potential (AP) velocity does not differ significantly for single iNGN neurons being spontaneously active and optogenetically activated APs of single iNGN neurons. Two-tailed Mann-Whitney test (Single cell: n =60 APs; Optogenetic single cell: n = 12APs). **c**, Response of a single fChRimson-iNGN neuron to several trains of light stimuli are shown. An increase in light intensity and light pulse duration leads to more APs elicited by one light pulse. For higher stimulation frequencies the latency between light on and AP peak are increasing. Left, middle: n =5, 10, 20, and 25 pulses; right: n = 5, 10, and 10 pulses for 0.5–5 Hz, respectively. Statistical analysis for 1^st^ APs at different frequencies: Shapiro-Wilk and Kolmogorov-Smirnov test for normality passed in all cases (alpha = 0.05) followed by ordinary one-way ANOVA with Tukey’s multiple comparisons test. Bottom plots: mean ± 95% confidence interval. **d**, The same behavior is observed for a different fChRimson-iNGN neuron (100 ms pulses, 46 µW/mm^2^). Lines represent one 4^th^ AP for 1 and 2 Hz. N = 5, 10, and 10 pulses for 0.5–2Hz respectively. **e**, Response of a single fChRimson-iNGN neuron to two trains of light stimuli at 2 Hz and 5 Hz are shown (50 ms pulses, 23 µW/mm^2^). Latencies between light on and AP peak are constant over the 2 Hz stimulation episode. For the 5 Hz episode, the first stimulus elicits an AP with similar latency as in the 2 Hz pulses, but all subsequent APs have increased latencies. In all plots, ns *P>0.05*, * *P≤0.05*, ** *P≤0.01*, **** *P≤0.0001*.

Next, we applied trains of light pulses that varied in duration, frequency, and intensity to single neurons expressing fChRimson (Fig. 4c). We found that a low irradiance (23 µW/mm^2^) with a pulse duration of 50 ms elicited one AP. By nearly doubling the irradiance (46 µW/mm^2^) but keeping the 50 ms pulse duration, two APs were consistently elicited up to 2 Hz. By further increasing the pulse duration to 100 ms, we were able to measure three elicited APs. A stepwise increase in number of APs can be achieved by modulating the light pulse intensity or duration. Notably, the latency, the time between light on and the peak of the AP, was about 10 ms longer for the low irradiance case compared to the high irradiance case. Delta latencies for subsequent spikes were defined as the time delay between the peaks of the previous and the current AP. Delta latencies measured at high light intensities with 50 ms and 100 ms pulse durations were similar but tended to be longer with increasing spike number. We confirmed the observations by measuring a second iNGN neuron that showed the same behavior while also firing a fourth AP in rare occasions (Fig. 4d). Another observation was that as the stimulation frequency increased, the latency also increased. By examining individual stimuli in the light train, we were able to confirm that the first light pulse does indeed elicit a response with the same latency as in the low-frequency case, but all subsequent spikes show an increased latency (Fig. 4e). Our observations could be explained by the light intensity used being close to the threshold of the rhodopsin leading to a slower increase in currents. Slowly depleting ion reserves in the cell would then lead to the change in latencies and number of fired APs with frequency and intensity. As shown here circuits and single neurons made with the SNAP method can be stimulated electrically and optically. Optogenetic stimulation and simultaneous MEA recordings provide a more natural environment for such measurements since they are non-invasive and ion concentrations are not controlled as opposed to patch clamp recordings. The method could thus help to characterize cells and optogenetic tools and to study information processing in circuits with high precision.

## Discussion

With our experimental approach, we show that endogenous electrical fields of a few neurons exhibit signatures of ephaptic coupling. Direct effects such as changes in AP velocity and synchronized activity can be measured longitudinally in a precise manner. Our results experimentally validate underlying theories and modeling studies on the effects of ephaptic coupling. These findings further support the notion that ephaptic coupling is a fundamental process in neural systems. Printed neuronal circuits in microfluidic devices with single cell resolution and persistent activity over weeks were key to obtaining these *in vitro* data. It is very likely that ephaptic coupling also plays an important role in *in vivo* systems, but this remains to be proven.

Our experimental platform serves as a tool to gain a more detailed understanding of the fundamental mechanisms underlying this phenomenon and allows the precise measurement of parameters due to purely endogenous fields. Future experiments should include myelinating cells such as oligodendrocytes. This could provide insight into the influence of myelin on ephaptic coupling. Since myelin is also thought to shield the effect of ephaptic coupling, the model could allow the study of the same circuits with high or low ephaptic coupling, allowing for the first time the design of experiments in which the effect of ephaptic coupling can be modulated. Modifying the setup to include simultaneous patch-clamp recordings may be of interest for future experiments. We have studied spontaneous neuronal activity as well as applied electrical and optogenetic stimulation, demonstrating the potential of SNAPs to address other fundamental questions in neuroscience. Incorporation of optogenetic activation could help to precisely control circuit activity at the single neuron and single spike level. Furthermore, the platform could be modified to serve as a disease model by introducing patient-derived neurons. In addition, the SNAPs presented show great sensitivity to synaptic antagonists, useful for drug screening applications.

## Methods

### Direct laser writing of microstructures

Microfluidic designs were created in Autodesk Inventor as a negative of the final structure. Custom designs were used except for the stomach-shaped structure inducing directionality which was adapted from Girardin et al.^42^ Microwells consisted of round structures with a diameter and height of 100 µm. Microchannels were 5µm high, keeping somas from migrating into channels^46^ and had a width of 20 µm (Y-shape) or 30 µm (directional structures). We used direct laser writing (DLW) for fabrication of the structures. The microfluidic designs were written on silicon substrates with four-inch diameters to quickly create many structures. Small, 25 x 25 mm^2^ fused silica samples were used to make and reproduce single structures. First, the substrates were cleaned, and an adhesion promoter (TI Prime, MicroChemicals GmbH) was applied via spin-coating. Then, resin IP-S (Nanoscribe GmbH & Co. KG) was applied. The designs were made using the Nanoscribe GT and the Nanoscribe QuantumX align systems in a layer-by-layer fashion. Lateral hatching distance of 0.3 µm and vertical slicing distance of 0.2 µm were used. To ensure a good print quality, while using a 25x magnification objective, the structures were split into cubes with 250 µm edge length. After the structures were exposed, they were developed using propylene glycol methyl ether acetate (PGMEA), rinsed with isopropanol, and blow-dried with nitrogen.

### Molding process

From the DLW-made master molds, microfluidics were prepared in a multistep casting process. First, master prints were silanized to lower the adhesion of the casting material. Vapor deposition of Trichloro(1H,1H,2H,2H-perfluorooctyl)silane (PFOCTS) was performed by placing samples on a tray over a few drops of PFOCTS in a desiccator. Pressure was reduced to -0.8 bar for 60 min and samples were baked at 90°C for 15 min subsequently. Base and curing agent of Polydimethylsiloxan (PDMS; SYLGARD® 184) was mixed thoroughly (10:1), and centrifuged until completely degassed and given on the master. After additional degassing in a vacuum chamber, the PDMS was cured at room temperature (RT) for 24 h and was subsequently silanized as described before. Epoxy resin (Resin L + Hardener L, R&G Faserverbundwerkstoffe GmbH) was prepared by mixing base and hardener 100:40, degassing via centrifugation, and directly casting onto the PDMS positive replica. Curing for 24 h at RT was followed by separating PDMS and epoxy and fully curing epoxy at 60°C for 5 h. After silanizing the epoxy negative replica, another PDMS positive replica was manufactured and silanized. To create molds for microfluidic fabrication, epoxy resin was given on the PDMS positive replica, gently degassed in a negative pressure chamber, pressed on a glass substrate, and taken off after curing at RT overnight. Excess material was removed and mold hardened at 60°C. A not completely closed frame of 100 µm thick transparency film was glued around the central structures. The molds were again silanized before microfluidic manufacturing.

### Microfluidics fabrication

PDMS was prepared as described previously, poured onto the glass-epoxy molds, and degassed in a vacuum chamber. A piece of transparency film was placed over the liquid PDMS in the frame and covered by a spacer made of a PDMS and a glass layer. With a clamp, the sandwich was pressed and placed in an oven to cure at 60°C for 30 min. After removing the microfluidics from the mold and frame we verified visually the structural integrity and that all microwells were open from the top. If not, we tried to get rid of the thin PDMS layer covering them by gentle scratching with a tweezer.

### Anti fouling surface coating

To prevent neurons extending on top of the microfluidic devices, we functionalized them with an anti-fouling surface coating as described previously^42,47^. Briefly, 13mg poly(allylamine hydrochloride) (PAAm; Sigma-Aldrich/Merck) were dissolved with 31.8 mg of potassium carbonate (Sigma-Aldrich/Merck) in 2.6 ml ultrapure water. The mixture was heated to boiling for improved dissolution and then allowed to cool rapidly. Separately, 11.2 mg of N-Succinimidyl 4-azido-2,3,5,6-tetrafluorobenzoate (ATFB-NHS; Iris Biotech) was dissolved in 4.135 mL of pure ethanol (EtOH) and protected from light, briefly ultrasonicated (∼10 s), and gradually added to the PAAm solution under continuous stirring with a magnetic stirrer. Stirring was performed for at least 3 hours to achieve a clear solution. If precipitation occurred, the mixture was discarded, and the process repeated. For further steps, the PAAm-ATFB solution was diluted to 0.1 mg/mL in a 2:3 mixture of HEPES/EtOH (10 mM of N-(2-Hydroxyethyl)piperazine-N′-(2-ethanesulfonic acid) (HEPES) in ultrapure water; Sigma-Aldrich/Merck). To functionalize the PDMS, all microfluidic devices were surface-activated in a plasma device (air, 18W power, 2 min; Diener electronic). The PAAm-ATFB solution was directly given on the microfluidics to cover them completely and left protected from light at RT for 30 min. After, PDMS was washed with HEPES/EtOH and ultrapure water. 10 mg/ml of polyvinylpyrrolidone (PVP; Sigma-Aldrich/Merck) in EtOH was added on top of the PDMS, excess solution was taken off, and everything blow dried with pressured air. The coating was then exposed to ultraviolet (UV) light in a cell culture hood for 5 min. To remove excess coating solutions, the PDMS was rinsed in methanol for 1 h, with the methanol being replaced every 15 min. Ultrasonication of microfluidics in fresh methanol for 5 min was done before washing with ultrapure water and storing of anti-fouling-coated microfluidics in ultrapure water at 4°C.

### Preparation of EM 1 cell line

EMX1 open reading frame was synthesized (Gen9, Inc), cloned into the doxycycline-inducible, puromycin-selectable lentiviral backbone pLIX_403 (Addgene #41395) using BP/LR Clonase (Thermo Fisher, Inc) following manufacturer protocols. Lentiviral particles were produced by co-transfecting pLIX403+EMX1, pMD2G (Addgene #12259) and psPAX2 (Addgene #12260) into HEK293T cells using polyethylenimine (PolyScience, Inc) as previously described^48^. iNGN cells were transduced with pLIX403+EMX1 lentiviral particles, then selected 48 hours post-transduction using 3µg/mL puromycin. For live cell microcopy, EMX1 cells were genetically modified to constitutively overexpress tdTomato.

### Stem cell culture and neuronal differentiation

Human neurons were generated by the overexpression of neurogenic transcription factors (TFs) in human induced pluripotent stem cells (hiPSCs). Induced neurogenin (iNGN) neurons were prepared as previously described^33,36,49^. By overexpression of TFs Neurogenin-1 and Neurogenin-2 either with or without EMX1 under a TetON inducible promoter system, hiPSCs differentiate within 4 days into post-mitotic neurons (either iNGN or EMX1). For live cell microcopy, iNGN cells were genetically modified to constitutively overexpress EGFP. After thawing, uninduced stem cells were cultured on Matrigel (Corning)-coated plates with mTeSR™1 medium (Stemcell Technologies), consisting of mTeSR™1 Basal Medium with mTeSR™1 Supplement and 1% penicillin-streptomycin (P/S; Thermo Fisher Scientific) added. Medium was exchanged every day. Cultures were passaged before reaching confluency by adding TrypLE for 3 min and centrifugation at 359 g for 4 min. After seeding, mTeSR™1 was supplemented with RHO/ROCK pathway inhibitor (ROCKi; Y-27632, Stemcell Technologies) for 24 h. Following at least two passages after thawing, differentiation of stem cells was induced on Matrigel-coated plates starting the day after passage by addition of 0.5 μg/ml doxycycline (Dox) (Sigma-Aldrich). Regular mycoplasma testing was performed on all cell cultures.

### Astrocyte culture

Astrocytes were prepared by expanding rat primary astrocytes (A1261301, Thermo Fisher Scientific) in cell culture flasks and used up to P4. Astrocyte media consisted of DMEM with 4.5 g/l d-glucose, pyruvate, N2 Supplement, 10% One Shot™ fetal bovine serum, and 1% P/S, all from Thermo Fisher Scientific. Astrocytes were cultured until almost confluent before reseeding in microfluidics.

### Single cell seeding

For preparation of defined neuronal circuits with single cell precision, a custom-built setup was used (see Supplementary Fig. 1b). It consists of a cell culture microscope (Evos XL Core, Thermo Fisher Scientific) with a microinjector (Narishige IM-11-2) mounted on a micromanipulator (Sutter Instruments MP-225) under a sterile cell culture hood. MEA chips (60 electrode MEAs, 60MEA200/30iR-Ti, Multi Channel Systems) or cover slips were treated with plasma (ambient air, 0.3 mbar, 50 W; Diener Electronics), incubated with Poly-D-Lysine (PDL, Merck) overnight at 37°C, 100% relative humidity (RH), and washed thrice with deionized sterile water. Microfluidics with anti-fouling coating were then placed on the MEA or cover slip with a drop of ultrapure water and aligned to the microelectrodes with a tweezer on a microscope under the cell culture hood. Substrates dried inside the hood to maintain sterility. Additionally, they were placed in a vacuum chamber for at least 20 min to evaporate all water and increase the attachment of the microfluidics. A rectangular frame cut from PDMS (∼3 mm wide and high frame with a ∼1x1 cm opening in the center) was placed around the microfluidics. After prewetting the microfluidics with ultrapure water and applying a short pulse of negative pressure to fill the microfluidic channels, the water was taken off and laminin (0.05 mg/ml; Sigma-Aldrich) added before incubating overnight at 37°C, 100% RH.

Rat primary astrocytes were reseeded on the microfluidics, usually the day before the single cell seeding. Astrocytes were detached with Accutase, centrifuged at 359 g for 4 min and resuspended in astrocyte media. 30,000 cells in 70 µl of media were given inside the frame on the microfluidics and incubated for 1 h before filling the MEA with astrocyte media and culturing until single cell seeding.

Just before single cell seeding, hiPSC-derived neurons at 5–6 dpi were detached with Accutase, centrifuged (359 g, 4 min) and resuspended in 1 ml of complete BrainPhys^TM^ medium. Complete BrainPhys^TM^ consisted of BrainPhys™ Neuronal Medium with 1% P/S, 2% NeuroCult™ SM1 Neuronal Supplement (all from STEMCELL Technologies), 1% N2 Supplement-A (Thermo Fisher Scientific), 20 ng/ml recombinant human BDNF, 20 ng/ml recombinant human GDNF (both from Peprotech), and 200 nM ascorbic acid (Sigma-Aldrich).

To investigate the best combination of supplements for increased survival of single neurons, we added either ROCKi or the CEPT cocktail (trans-ISRIB, 0.7µM, Cayman Chemical; chroman 1, 50nM, MedchemExpress; emricasan, 5µM, Cayman Chemical; 0.1% Polyamine Supplement, Sigma-Aldrich) as reported by Chen et al.^38^ to the culturing medium. Typically, cytarabine (AraC) is added to differentiating stem cell cultures after induction and just before reseeding to eliminate undifferentiated and still dividing cells. Since AraC has been reported to be neurotoxic and to cause DNA damage^50^, we tested omitting it. By adding CEPT without AraC treatment, the survival rate increased to about 62.5% of cells 28 days after seeding (Supplementary Fig. 1c).

Single cell seeding was performed after detachment and resuspension of neurons in supplemented (-AraC +CEPT for best performance) BrainPhys^TM^ medium. Astrocyte medium was aspirated from the samples, washed once with DPBS+/+ (Thermo Fisher Scientific) and supplemented BrainPhys^TM^ was added inside the PDMS frame. A small amount of BrainPhys^TM^ was added in the surrounding area outside the frame so that the two media reservoirs were not connected. A drop of resuspended neurons was added to the surrounding media, with the frame preventing cells from flowing onto the microfluidics. Using the custom setup, single neurons were picked up with a glass pipette (VESbl-12-0-0-55, BioMedical Instruments) from the outside of the frame and then transferred to the inside and placed in a microwell. The correct placement of the cell in the microwell was visually confirmed before repeating the procedure until all microwells contained a neuron. The media outside the edge was aspirated, the area gently washed, and the MEA covered with a lid and placed in the incubator. After the cells had attached, the MEAs were filled with BrainPhys^TM^. Five days after the initial seeding of single neurons into the microwells, a second round of seeding was performed to maximize the number of complete circuits. The procedure was repeated and any microwells where the previously placed neuron had been lost were reseeded. The old media was collected from the MEAs and enough media was left in the MEA for the seeding procedure (PDMS edge filled with media, small amount in the surrounding area). After placing the neurons, the environment was aspirated, washed, and the MEAs were left in the incubator for attachment. Later, media (1:1 old media and fresh complete BrainPhys^TM^ with CEPT) was added (Supplementary Fig. 1a).

As a control, populations of neurons were seeded into the microfluidic structures. This was done in parallel with single cell seeding. After aspiration of the astrocyte medium and washing of the MEA, 15,000 resuspended neurons in 70 µl of complete BrainPhys^TM^ with CEPT were added within the PDMS edge of the microfluidics. A second round of seeding was performed with 15,000 neurons in 12 µl of media to fill all microwells with at least five neurons, as visually confirmed under the microscope. After leaving the MEAs in the incubator for attachment, they were filled with complete BrainPhys^TM^ with CEPT.

For testing optogenetic stimulation of single neurons, we seeded single cells of the fChRimson-iNGN cell line in microfluidics. The fChRimson-iNGN cell line contained a double-floxed inverse open reading frame (DIO) of the fChRimson-EYFP gene^45^. The expression of fChRimson was induced by transfecting the cells with Cre recombinase mRNA, thereby, the fChRimson gene was in frame. This is crucial, since stem cells often lose the expression of transgenes, especially optogenes. The DIO system can be used to delay the expression of the optogene until after the neuronal differentiation has been started by overexpression of TFs^51^. The fChRimson is a fast variant of the red-shifted channelrhodopsin ChRimson with an absorption maximum at around 590nm wavelength of light ^45,52^. The mRNA was prepared by amplifying the Cre DNA via PCR using the following primers: T7-Cre-forward: 5’-GCTAATACGACTCACTATAGGGACAGGCCACCATGGCCAATTTACTGA-3’ and T7-Cre-reverse: 5’-TCATTACGGTCCATCGCCATCTTCCAGCAGGCGCACCATT-3’ from the pCAG-Cre:GFP plasmid. pCAG-Cre:GFP was a gift from Connie Cepko (Addgene plasmid # 13776)^53^.

The HiScribe^®^ T7 ARCA mRNA Kit (with tailing) (New England Biolabs) was used to create mRNA and subsequently cleaned up with the Monarch^®^ Spin RNA Cleanup Kit (50 μg) (New England Biolabs). After transfecting the cells with Cre-mRNA using Lipofectamine™ MessengerMAX™ (Thermo Fisher Scientific) according to the manufacturers protocol at 0 dpi, the expression cassette in the cells’ gene was flipped and fChRimson was expressed. Cells were treated and seeded as described above.

### Electrophysiology experiments

Regular MEA recordings were performed on a MEA2100-Lite-System (Multi Channel Systems) and imaging using a Keyence BZ-X810 using a 20x Keyence BZ-PF20LP objective. Recordings were performed for at least 3 min, usually 5–10 min at following dpi (± 1 day) for Y-structures: dpi 13, 22, 27, 33, 39, 49, and for directional structures: dpi 14, 18, 21, 24, 27, 31, 33, 36, 49, 54, 61, 67, 76. Half media was exchanged weekly and fresh complete BrainPhys^TM^ medium added.

Electrical stimulation was performed using the MC_Rack software (Multi Channel Systems). After recording the baseline activity of neuronal circuits, a train of stimulation pulses (30 biphasic pulses consisting of -15 µA for 100 µs followed by 15 µA for 100 µs with interpulse interval of 1800 ms) was applied to the left and right microchannels of the directional structures while recording the activity.

Optical stimulation of fChRimson-iNGN cells was done by placing the MEA headstage under an upright microscope (Scientifica SliceScope) with a 10x objective (Olympus Plan N 10x/0.25). A LED light source (CoolLED pE-800) was connected to the microscope. A protocol for the light stimulation was prepared with the Clampex 11.1. software that controlled the light source. Stimulation time stamps and MEA electrophysiology data were recorded with MC_Rack. The same stimulation protocol was applied with two different irradiances: 23 µW/mm^2^ and 46 µW/mm^2^. Low irradiance levels were used to avoid a possible cytotoxic effect to single neurons and to prevent stimulation artifacts in the MEA electrodes. Stimulation was performed through the lid of the MEA to keep it sterile. Diameter of the light spot was 2 mm and centered on the electrode area, covering it fully. The protocol consisted of trains of 50 ms light pulses with 0.5 Hz, 1 Hz, 2 Hz, and 5 Hz followed by 100 ms light pulses with 0.5 Hz, 1 Hz, and 2 Hz.

To block transmission between neurons through excitatory or electrical synapses, antagonists were added to the cultures. Excitatory NMDA and AMPA synapses were blocked by adding DL-2-Amino-5-phosphonopentanoic acid (AP5; Tocris) and 2,3-Dioxo-6-nitro-1,2,3,4-tetrahydrobenzo[*f*]quinoxaline-7-sulfonamide disodium salt (NBQX; Tocris), while electrical synapses (gap junctions) were blocked using carbenoxolone (CBX; Sigma-Aldrich). First, a baseline of activity was recorded of all samples. If stimulation was part of the overall experiment, a stimulation protocol was also applied during all conditions. Part of the media was removed and collected for later use so that the cultures were covered still. NBQX (10 µM final concentration) together with AP5 (50 µM final concentration) were added to the media and incubated for 30 min. Then activity was recorded again. After aspiration of the media and washing of samples (1x DPBS+/+, 2x fresh BrainPhys^TM^), the collected old media was added to the culture and incubated for 1 h before recording the activity again. CBX (100 µM final concentration) was added in a second step using the same protocol.

### Data analysis of electrophysiological recordings

Electrophysiology data was processed with custom-made python scripts using the SpikeInterface package^54^. Preprocessing of data consisted of a second order high pass Butterworth filter with a cutoff at 100 Hz and a subsequent global common median referencing step. In SNAP recordings, spike sorting algorithms were not able to separate spike trains of single neurons from the activity reliably (Supplementary Fig. 2d), likely because of the densely packed APs^55^. Therefore, we extracted all peaks in the traces of all electrodes recording from the circuit using the following criteria: negative signs only, detection threshold: 5 x median absolute deviations, exclude_sweep_ms: 0.2. Detected peaks and their timestamps were used as spike trains for further analysis of network activity.

AP velocities were extracted from preprocessed raw traces using a custom-made python script. Spikes in the recording were chosen at random and an estimate of the timepoint in the range of one ms before the AP was recorded was fed into the algorithm. The algorithm then calculated the standard deviation of the signal during the 10 ms before and extracted the timepoint when the trace rose above the threshold of five times the calculated standard deviation. This was done in all electrodes and timepoints of the threshold crossings were used to calculate the propagation velocities. Distances were known from the microfluidic designs. A minimum of ten APs per condition, sample and timepoint were extracted. If during the 10 ms before the AP there was another AP visible in the trace, the AP was rejected for measurement since the calculation of standard deviation would be distorted.

Interspike interval (ISI) histograms were prepared by binning all time intervals between two consecutive spikes extracted in an electrode (1 ms bin size). The Kullback-Leibler divergence^40^ of two probability distributions P(x) and Q(x) is defined as:

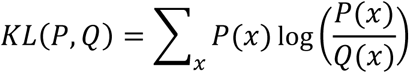

and gives a measure of how much two distributions differ. This measure was applied to ISI histograms, comparing histograms of baseline activity and activity under excitatory blockers. All electrodes in a motif with single neurons or populations of neurons were evaluated separately. To avoid artifacts in the calculation of the KL divergence because of sparse histograms we only included histograms with more than 400 ISI values in the analysis.

As a measure of synchronicity in a circuit we used the Spike-contrast algorithm^43^ via its implementation in the Elephant python package^56^. Burst extraction was performed with a MaxInterval method^57^ with following parameters: minimum interburst interval (IBI): 200 ms, minimum burst duration: 1 ms, minimum number of spikes in burst: 3, maximum ISI in a burst: 10 ms. For extraction of synchronized network bursts, we only accepted bursts that were present in all electrodes in a window of 5 ms. To extract the participation ratio (PR) of a circuit motif, we calculated the matrix of correlation coefficients of all binned spike trains in a circuit (5 ms bins). The PR is defined by:

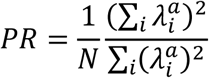

where 𝜆^𝑎^ are the eigenvalues of the covariance matrix of the activation of N units^44^. This ratio 0 ≤ PR ≤ 1 gives an intuition of the dimensionality of the activity in a network. If the neurons’ activity is more decoupled, the dimensionality and the PR are large, if the neurons’ activity is more coupled and correlated, the dimensionality and PR are smaller, meaning more neurons participate in the common activity.

For the analysis of responses to optogenetic stimulation, we extracted the AP peaks in a window of 150 ms after the light-on timepoint. To exclude artifacts, we only accepted peaks with an amplitude greater than 50 µV.

### Immunofluorescence

Neuron cultures were fixed with 4% paraformaldehyde solution (Thermo Fisher Scientific) for 15 min at RT. After washing with PBS, permeabilization with 0.2% Triton X-100 (Sigma-Aldrich/Merck) in PBS and 30 min of blocking with 5% donkey serum (Sigma-Aldrich/Merck) in PBS were performed at RT. Primary antibodies were added overnight at 4°C (anti-GFP, 1:200, A10262, Thermo Fisher Scientific; anti-beta-III-tubulin linked to eFluor^TM^ 570, 1:200, 41-4510-80, Invitrogen). Washing was done 3x with 0.1% Triton X-100 in PBS. Secondary antibody was added in blocking solution for 2 h at RT (goat anti-chicken Alexa Fluor 488, 1:500, A11039, Thermo Fisher Scientific). After washing 3x with 0.1% Triton X-100 in PBS, Hoechst stain (H3570, Thermo Fisher Scientific) was added for 5 min at RT (1:2,000 in PBS). After washing 3x with PBS, samples were mounted in ProLong^TM^ Diamond Antifade Mountant (Thermo Fisher Scientific). Imaging was done on an Echo Revolve microscope.

### Statistical analysis

Statistical analysis was performed in GraphPad Prism version 10.4.1 for Windows, GraphPad Software, Boston, Massachusetts USA, www.graphpad.com. Plotting of data was done in GraphPad Prism and with custom-made python scripts. Statistical tests are mentioned in the figure captions where applicable with corresponding number of values. Survival rates were extracted by evaluating the number of neurons visible in the fluorescence microscopy images per circuit.

## Acknowledgements

We thank Dr. Sarah Kunze and Dr. Kirsten Harmening for technical assistance. We also thank Prof. Dr. Andreas Offenhäusser, Prof. Dr. Heinz Beck, Prof. Dr. Dr. Florian Mormann, Dr. Johannes Kirchner, Dr. Christoph Miehl and Dr. Robert Prior for their comments and the discussions. We are grateful for technical assistance and comments by the Gene Editing Core Facility at the University Hospital Bonn with Dr. Simon Schneider and Angela Egert. JS acknowledges support from the Joachim Herz Foundation. HG acknowledges support from the Studienstiftung des deutschen Volkes. VB acknowledges support from the Deutsche Forschungsgemeinschaft (DFG – project number 531985111), the Volkswagen Foundation (Freigeist – A110720), the German Federal Ministry for Economic Affairs and Climate Action through the German Space Agency at DLR in the framework of SANSRETINA (grants: 50WB2516), the Pro Retina Foundation and the Paul Ehrlich Foundation. VB is also supported by the Cluster of Excellence – ImmunoSensation2 EXC-2151-390873048 at the University of Bonn.

## Author contributions

Conceptualization: J.S. and V.B.; Methodology: J.S. and V.B.; Software: J.S.; Validation: J.S.; Formal analysis: J.S.; Investigation: J.S., R.H., D.W., H.G., E.P., J.P., K.S., A.H.M.N. Resources: J.S., W.P. and V.B.; Data Curation: J.S.; Writing - Original Draft: J.S.; Writing - Review & Editing: all authors; Visualization: J.S., R.H. and E.P.; Supervision: W.P. and V.B.; Project administration: J.S. and V.B.; Funding acquisition: J.S. and V.B.

## Competing interests

A.H.M.N. and V.B. are inventors on patents filed by the Presidents and Fellows of Harvard College. A.H.M.N. is a co-founder of and has equity in GC Therapeutics, Inc.

## Additional Information

Supplementary Information is available for this paper.

Correspondence and material requests should be addressed to Prof. Dr. Volker Busskamp.

During the preparation of this work the authors used DeepL-Write Pro to improve the readability and language of the manuscript. After using the tool, the authors have reviewed and edited the content as needed and they take full responsibility for the content of the published article.

## Data availability

The data generated and analyzed during the current study are available from the corresponding author on reasonable request.

## Code availability

Custom computer code used for analysis of the data is available from the corresponding author on reasonable request.

---

**Supplementary Fig. 1.**
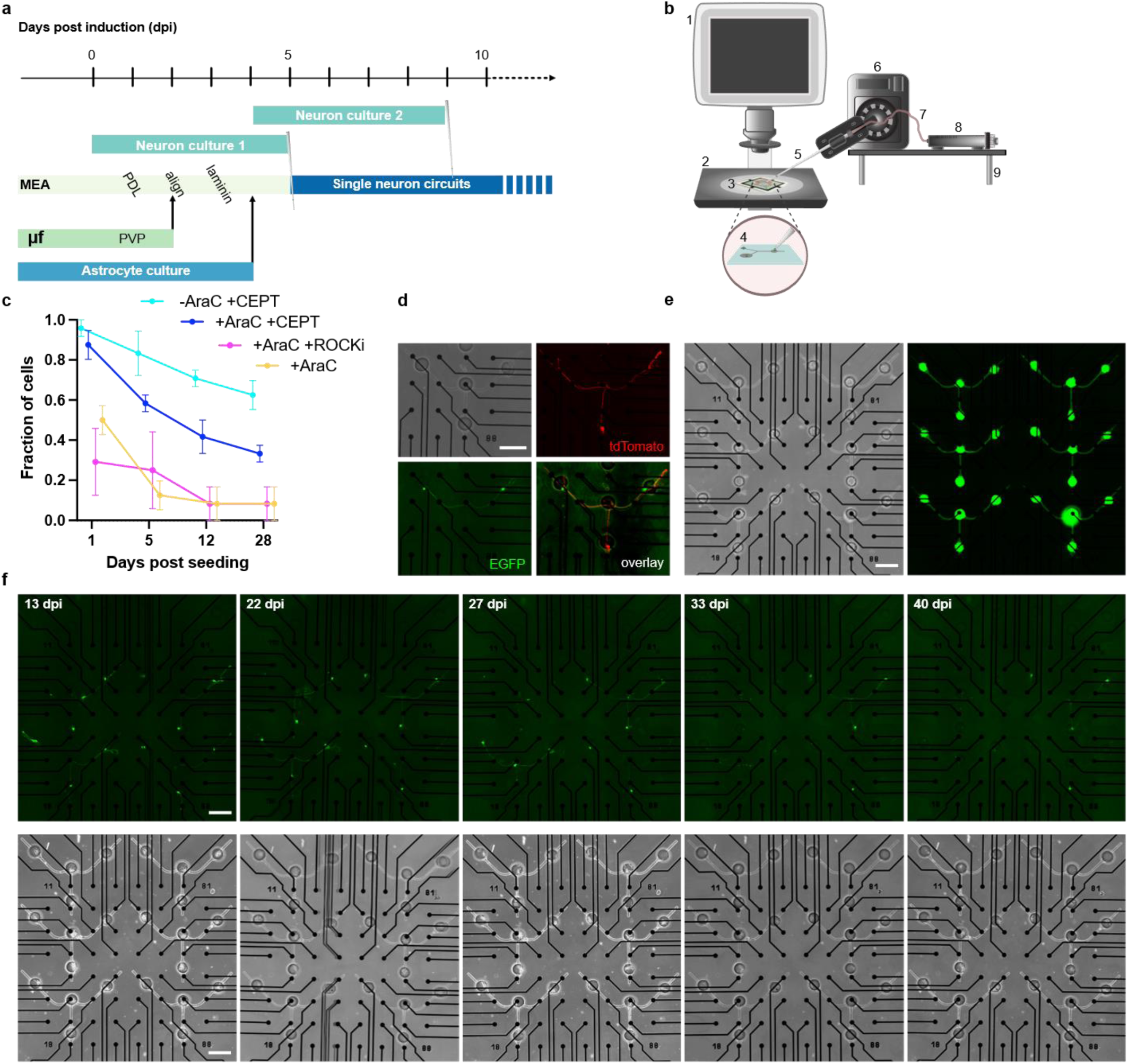
: Preparation and time development of single neuron circuits. a, Overview of the single neuron circuit preparation protocol. Single cell seedings are marked with a glass pipette icon. µf: microfluidics, MEA: multielectrode array, PVP: polyvinylpyrrolidone functionalization, PDL: Poly-D-Lysin coating. **b**, Cell placement setup installed in a sterile cell culture hood and consisting of a microscope (1) and stage (2) on which a MEA with microfluidics (3) is placed for seeding (4). A micropipette (5) is steered by a micromanipulator (6). A silicon tube (7) connects the manual microinjector (8) to the pipette. All cell placement equipment is mounted on a custom-build and adjustable platform (9). **c**, Comparison of cell survival under different preparation conditions. Fraction of surviving iNGN neurons is shown for different days post seeding. Adding CEPT cocktail and no AraC showed the best cell survival. N = 3 per condition and timepoint. **d**, Representative neuronal circuit motif consisting of two cell types: two iNGN (upper microwells) and one EMX1 (lower microwell) neurons connected through microchannels within a microfluidic device. Fluorescence images of tdTomato constitutively expressed by EMX1 neurons, and constitutively expressed EGFP by iNGNs. Scale bar, 200 µm. **e**, Representative MEA with populations of iNGN neurons seeded in microfluidics at 22 dpi (EGFP and brightfield channels). **f**, Development of four Y-shaped motifs with 3xiNGN on one MEA is shown across time (EGFP and brightfield channels). All scale bars, 200 µm.

**Supplementary Fig. 2.**
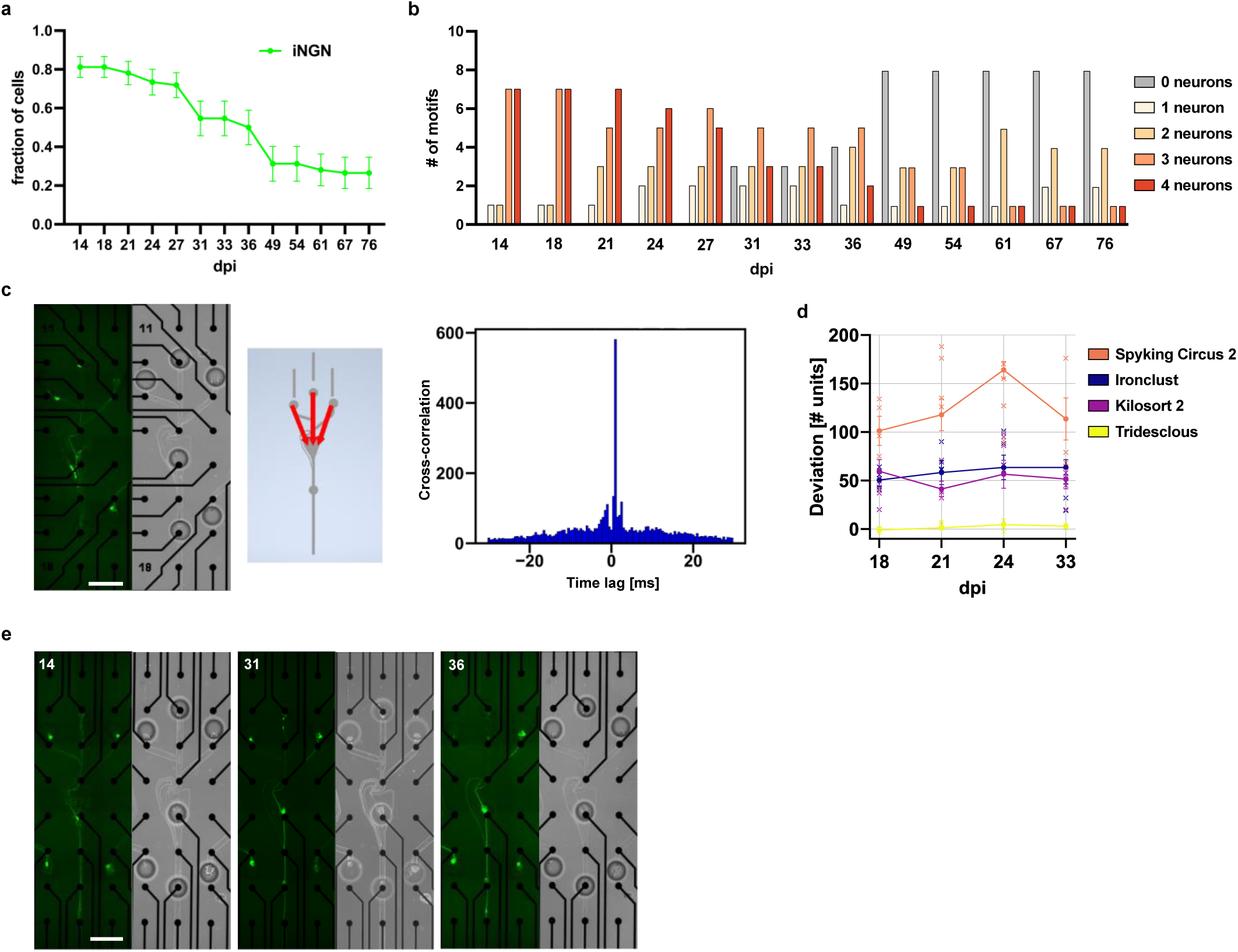
: Survival and functional analysis of directional circuits. **a**, Fraction of viable cells at several days post induction (dpi) for directional 4-iNGN circuit motifs (mean±s.e.m., iNGN: n = 16 circuits). **b**, Number of circuit motifs containing respective number of iNGN neurons per dpi. 4 samples with 4 motifs each were monitored. **c**, Representative analysis of signal propagation in a directional circuit of 4 iNGN neurons. Left: Fluorescence (constitutively expressed EGFP) and brightfield images of the circuit. Middle: Schematic of the microfluidic design overlayed with arrows indicating the direction of AP propagation as extracted by cross-correlation analysis of the signals measured in the electrodes of the respective microfluidic channels. Right: Representative cross-correlation analysis of signals in two electrodes measured in the circuit. **d**, Performance of different spike sorting algorithms on directional circuits. Four algorithms were compared by extracting the number of units from four MEAs on four different dpi. The true number of units as observed in microscopy images was subtracted from the number of units found by the algorithms to get a deviation value. mean±s.e.m. **e**, Fluorescence (constitutively expressed EGFP) and brightfield images of the circuit described in Fig. 3e with respective dpi mentioned. All scale bars, 200 µm.

**Supplementary video 1: Single cell seeding process.** Cells are picked up by a glass micropipette from an area where a cell pool is kept and transferred to the electrode area of the multielectrode array with the microfluidic structure on top. A micromanipulator allows precise control of the micropipette in all three directions. A microinjector is used to pick up and release the cells. Speed of the video was tripled.

